# C5aR Expression in Kidney Tubules, Macrophages and Fibrosis

**DOI:** 10.1101/2023.12.29.573612

**Authors:** Carolyn Dunlap, Niky Zhao, Linda S. Ertl, Thomas J. Schall, Kathleen M. C. Sullivan

## Abstract

The anaphylatoxin C5a and its receptor C5aR (CD88) are complement pathway effectors implicated in renal diseases, including ANCA-associated vasculitis. We investigated the kidney expression of C5aR and a second C5a receptor C5L2 by using immunohistochemistry and in situ hybridization on formalin-fixed, paraffin-embedded human and mouse kidney. C5aR was detected on interstitial macrophages and in multiple tubular regions, both distal and proximal; C5L2 had a similar expression pattern. The 5/6 nephrectomy model of chronic kidney injury exhibited increased C5aR expression by infiltrating cells within the fibrotic regions. Functional assessment of myeloid C5aR in vitro revealed that C5a induced the expression of chemokines and remodeling factors by macrophages, including CCL-3/-4/-7,-20, MMP-1/-3/-8/-12, and F3, and promoted survival by blocking neutrophil apoptosis. C5a activity was C5aR dependent, as demonstrated by reversal with the C5aR inhibitor avacopan. Collectively, these results suggest that myeloid C5aR may induce excessive inflammation in the kidney via immune cell recruitment, extracellular matrix destruction, and remodeling, resulting in fibrotic tissue deposition.

## INTRODUCTION

Complement is a network of over 30 soluble and membrane-associated proteins that rapidly initiate enzymatic cascades in response to microbial or other stimuli, resulting in the formation of anaphylatoxins C3a and C5a, which mediate inflammatory responses, and the membrane attack complex C5b-9, which induces cell lysis (1). The peptide C5a is produced from the cleavage of complement factor C5 and is a ligand for the G-protein coupled receptor C5aR (C5aR1 or CD88) and the atypical GPCR C5L2 (C5aR2 or CD77) (2). C5aR is expressed in circulating myeloid cells and contributes to inflammatory responses via activation of these cell types (3). Genetic deletion and pharmacological inhibition studies in disease models have implicated C5aR in the pathogenesis of multiple diseases including ANCA-associated vasculitis, kidney fibrosis, infection, and cancer (4–7). C5L2 function is less well understood (8), but genetic deletion of C5L2 has shown opposing effects to C5aR in disease models, suggesting it may be involved in negative regulation of C5aR in some contexts, including ANCA vasculitis (5, 7, 9).

A potent antagonist of C5aR, avacopan, showed superior ability to induce and maintain remission in ANCA-associated vasculitis patients with renal involvement, compared to high dose oral prednisone in a phase III trial (10). Avacopan significantly improved patient kidney function by reducing proteinuria and increasing the estimated glomerular filtration rate, reproducing similar activity observed in phase II trials (11). Avacopan may contribute to disease remission by blocking neutrophil activation and tissue migration induced by C5a, which results from anti-neutrophil antibodies or other stimuli that activate complement (12, 13). The continued improvement in renal function after induction of disease remission (10) suggested that C5aR-expressing cells within the kidney may contribute to avacopan disease-modulating activity.

The expression pattern of C5aR protein and mRNA in normal and diseased kidney has been examined in numerous studies that have produced conflicting results. C5aR has been reported to be expressed (14–16) or not expressed (17–21) on interstitial cells within the human, rat, and mouse kidney, which through serial section staining have been tentatively identified in some cases as macrophages (14, 16). C5aR expression has also been reported on normal human and mouse kidney tubules (18, 20, 22–24); however, the tubule region expressing C5aR has been variously characterized as proximal (24), or proximal, distal, and other (18, 20), or not specified (22, 23). Tubule expression is controversial, as several groups have detected no C5aR in normal human or rodent kidney tubules (15–17, 21). C5aR-expressing tubules have been reported in several kidney diseases and disease models, in some cases by groups that detected no tubule expression in normal samples: diabetes (19), IgA nephropathy (17), sepsis (16) and ischemia/reperfusion (21, 23). A single report on C5aR in ANCA patient kidney biopsies observed a significant decrease in tubule expression compared to normal (20). There has been similar variability in reported expression of C5L2 in mouse and human kidney: widespread (20), in tubular subsets (18), or in both tubular and interstitial cells (25).

Given the apparent contradictory findings within the literature, our initial aim was to better understand C5aR expression in normal human and mouse kidney by examining protein and transcript with validated methods and reagents. As an extension of these studies, myeloid C5aR expression and function was investigated in vitro, to identify kidney-relevant roles of C5aR in these cell types.

## MATERIALS AND METHODS

### Immunohistochemistry

FFPE blocks of adult human non-diseased kidney samples were obtained from Cureline donors COR/MED (female age 63), LL (male age 60), kidney cortex C (male age 41), and kidney B (male age 60), all of which were collected within 1 hr postmortem. Sections of 5 µm were cut from the FFPE blocks onto Superfrost plus slides and baked at 60°C for 1 hour, then deparaffinized in 3 steps of xylene (VWR 89370-090) followed by graded steps of ethanol (3 steps at 100%, 3 steps at 95%, 2 steps at 70%, and 2 steps at 50% with reagent alcohol (VWR BDH1156-4LP)) into water. Heat induced epitope retrieval (HIER) was conducted using target retrieval solution pH 9 (Dako S236784-2, 10x concentrate diluted to 1x with distilled water and pre-heated in the microwave) for 20 minutes at 95°C, followed by a 20 minute hold at room temperature (RT) and 2-3 washes for 5 minutes total with distilled/deionized water. Samples were then treated with BLOXALL Endogenous Peroxidase and Alkaline Phosphatase Blocking Solution (Vector Laboratories SP-6000) for 10 minutes, washed in distilled water and blocked for 30 minutes with 5% normal goat serum (Jackson ImmunoResearch 005-000-121) in PBS-T (PBS (VWR 45000-436), 0.5% bovine albumin serum (Sigma Aldrich A7979), and 0.5% Triton X-100 (Sigma Aldrich 93443)).

The anti-human C5aR monoclonal antibodies used for IHC were as follows: S5/1 (Hycult Biotech Cat# HM2094, RRID:AB_533292), P12/1 (Bio-Rad Cat# MCA2059, RRID:AB_566906), 8D6 (Thermo Fisher Scientific Cat# MA1-70060, RRID:AB_1073841). Primary antibodies or their respective mouse (R and D Systems Cat# MAB003, RRID:AB_357345) or rat (BD Biosciences Cat# 559478, RRID:AB_10056899) isotype controls were added to the samples at 2 µg/ml for overnight incubation at 4°C. Samples were washed four times with 1% goat serum PBS-T for 5 minutes each, then incubated with 1:1000 diluted, HRP-conjugated goat anti-rat or anti-mouse secondary antibodies as appropriate (Jackson ImmunoResearch Labs Cat# 112-035-167, RRID:AB_2338139, and Cat# 115-035-062, RRID:AB_2338504) in 1% PBS-T for 1 hour at RT, after which DAB substrate (Vector Laboratories SK-4105) was applied for 5 minutes. Samples were counterstained with Mayer’s hematoxylin (Sigma MHS32) for 30-60 seconds, then dehydrated by moving slides through the following solutions in order, 3 repetitions each: 95% ethanol, 100% ethanol, and xylenes. To mount slides, 1-2 drops of Fisher Chemical Permount Mounting Medium (ThermoFisher SP15-100) were added to the sample and topped with a coverslip, then allowed to harden for 1-2 hr prior to imaging on a Nikon Ti2 microscope with an Ri2 camera.

### In situ hybridization and dual ISH/IHC

The adult human non-diseased kidney FFPE cores used for ISH analysis included two of the Cureline donors used for IHC (CORT/MED and LL) and core needle biopsies from Biochain (T2234142-D03), donor lots as follows: C611197 (male age 44), C611198 (male age 65), and C611199 (male age 51). The Cureline donor LL sample was used for dual ISH/IHC staining. All samples were confirmed to have high levels of the positive control PPIB transcript by ISH.

The FFPE blocks were cut at 5 µm and baked on Superfrost plus slides for 1 hour at 60°C. ISH was performed using the RNAscope 2.5 HD detection (RED) assay with the following probes from ACD biosciences: anti-sense dapB (bacterial gene), human C5aR, C5L2, and PPIB, and sense human C5aR. Manufacturer’s instructions for RNAscope protocol were followed, with all kidney samples subjected to 15 minutes of target retrieval and 30 minutes of protease treatment, as determined by optimizing pretreatment conditions using the housekeeping gene PPIB. All sections were imaged using a Nikon Ti2 microscope and Ri2 camera.

For dual ISH/IHC for cell markers, samples were first processed for ISH with anti-sense C5aR probe. The IHC protocol was performed as described, excluding the HIER antigen retrieval process, using one of the following: affinity purified rabbit polyclonal anti-CALB1 or anti-SLC13A3 (Sigma-Aldrich Cat# HPA023099, RRID:AB_1845859 and Cat# HPA014736, RRID:AB_1856970), or anti-CD68 (Abcam Cat# ab192847).

### Mouse human C5aR knock in 5/6 nephrectomy study

All study procedures were carried out in accordance with the study protocol approved by the ChemoCentryx Institutional Animal Care and Use Committee. Human C5aR knock-in mice were generated in the C57BL/6 background by ChemoCentryx (7) and maintained as a colony in accordance with guidelines described in the Guide and Use of Laboratory Animals of the National Research Council. Nephrectomies were performed on 12 human C5aR knock-in male mice aged 7-10 weeks by a surgeon contracted from Charles River Laboratories. The right kidney was completely removed, with renal blood vessels and ureter ligated, and the left kidney was ligated around each pole at the one-third position and excised, leaving one-third of the left kidney remaining. The incision was closed in two layers, and the wound clips were removed 7-10 days post surgery. During the surgical procedure mice were anesthetized with isofluorane and kept on a circulating water blanket until fully recovered and returned to cages. Mice were monitored for weight changes and signs of morbidity. Torbugesic was given prior to surgery and additional analgesic afterwards when signs of pain or discomfort were observed. Post surgery all mice lost weight and were given DietGel and HydroGel; 10 mice recovered and gained weight.

Kidneys were harvested 6 weeks post surgery from these and age-matched male hC5aR KI mice (n=5) that did not undergo surgery. Kidneys were weighed and immediately placed in a formalin jar (VWR prefilled 10% neutral buffered formalin 16004-115). After 24 hours of fixation the kidneys were transferred to 70% ethanol for another 24 hours. Samples were processed and cut according the IHC or ISH protocol used for human samples. For IHC the primary anti-C5aR antibody 8D6 was followed by incubation with a biotin-labelled, goat anti-rat secondary antibody (absorbed for mouse, Jackson ImmunoResearch, diluted 1:1000), then avidin-HRP (Vector Laboratories, diluted 1:1000).

Trichrome staining (Gomori One-Step, Aniline Blue Stain Kit, Fisher Scientific, 9176A) was performed by deparaffinizing sections and holding them mordant in Bouin fluid for 1 hour at 60°C in a water bath. Sections were counterstained using Weigert’s iron hematoxylin for 10 minutes, and rinsed in water for an additional 10 minutes. Next the Gomori One Step, Aniline Blue stain was applied for 20 minutes, followed directly by differentiation using 0.5% acetic acid for 2 minutes and rinsing with distilled water. Sections were dehydrated using two changes of 95% ethanol and 100% ethanol each, cleared with 3 steps of xylene and coverslipped using permount.

### In vitro assays

#### Human leukocyte isolation and staining

Human leukocytes were isolated from healthy donor blood enriched with mononuclear cells in Leukoreduction System (LRS) chambers (Stanford Blood Center, Stanford, CA). The chamber content (∼10 ml) was transferred to a 50 mL sterile centrifuge tube, diluted with PBS to 35 ml, then layered over 14 mL Ficoll-Paque Plus in SepMateTM-50 tubes (STEMCELL Technologies 85450). The tubes were centrifuged at 2800 revolutions per minute (rpm) for 10 minutes at room temperature (RT). The PBMC layer was transferred to a 50 mL conical tube and red blood cells lysed with Pharm Lyse buffer (BD 555899) for 10 minutes at RT. The cells were immediately processed as described below for each assay.

For flow cytometry analysis, whole blood was directly stained with the following antibodies from BioLegend: C5aR (Cat# 344304, RRID:AB_2067175); C5L2, (Cat# 342404, RRID:AB_2247831); CD16 (Cat# 302039, RRID:AB_2561354); CD45 (Cat# 368521, RRID:AB_2687374); CCR3 (Cat# 310719, RRID:AB_2571958); Siglec-8 (Cat# 347107, RRID:AB_2629715); HLA-DR (Cat# 307641, RRID:AB_2561360); CD14 (Cat# 301837, RRID:AB_11218986). Each immune cell type was identified using the following gating scheme on a BD LSRFortessa™: SSC-A^HI^CD16+ for neutrophils; SSC^LO^CD45^DIM^HLA-DR-CD123+ for basophils; CD16-CCR3+Siglec8+ for eosinophils; and CD45+CD14+ for monocytes.

#### Macrophage differentiation and function

Monocytes were isolated from PBMCs using human CD14+ MicroBeads (MACS Miltenyi Biotech, 130-045-201) and an autoMACS®Pro Separator according to manufacturer’s specifications. The monocytes were resuspended in RPMI-1640 (Corning 10-041-CM) with 10% FBS and 1X penicillin-streptomycin (100X, Corning 30-002-CI) at 1×10^6^ cells/ml in T-175 flasks with 100 ng/ml of either GM-CSF or CSF-1 (PeproTech 300-23 or 300-25). Monocytes were incubated at 37°C/5% CO_2_ for up to 6 days to differentiate into macrophages, replacing 50% volume on day 3 with fresh media including cytokines. Macrophages were then resuspended in RPMI-1640 with 10% FBS at 1×10^6^ cells/ml, seeded at 2 ml per well in 12 well flat-bottom plates and treated with C5a (PeproTech, 300-70) or avacopan for 72 hrs. Supernatants were collected and analyzed for the presence of secreted factors using Luminex human Discovery Assays (R&D Systems): LXSAHM-02 (MMP-8), −04 (MMP-2, S100A12, MPO), and −29 (MMP-9, MIP-1α/CCL3, GRO-α, TNF, MIP-1β/CCL4, MIP-3α/CCL20, IFNɣ, IL-6, IL-1α, IL-1β, MDC, IL-8, Fractalkine, eotaxin, VEGF, IFNβ) and DLAP00 (Human LAP (TGF-β1)). To measure cell surface F3 levels, macrophages were stained with APC-conjugated antibody clone NY2 (BioLegend Cat# 365205, RRID:AB_2564567) and subjected to flow cytometry.

#### Neutrophil survival

PBMCs were washed twice and adjusted to 2×10^6^ cells/ml in HBSS plus 5% FBS (Sigma-Aldrich F2442). To induce apoptosis, the cells were incubated in buffer with 0.025% H_2_O_2_ or 500 ng/ml anti-CD95 (Sigma-Aldrich 05-201) for 2 hr, with or without C5a or avacopan. The cells were then stained with FITC Annexin V Apoptosis Detection Kit with 7-AAD (BioLegend 640922) following the manufacturer’s instructions and subjected to flow cytometry. Both apoptosis assays were performed with similar results in 4 donors.

## RESULTS

### C5aR was expressed in non-diseased human kidney tubule and macrophages by in situ hybridization and immunohistochemistry

Three anti-human C5aR monoclonal antibodies, S5/1, P12/1, and 8D6, were evaluated on leukocytes from human blood by both flow cytometry and immunohistochemistry (IHC). All clones readily detected C5aR on neutrophils as expected (3, 26), when compared to isotype controls (Supplemental Figure 1). Despite similar staining of leukocytes, the clones had divergent expression patterns by IHC in sections from formalin-fixed, paraffin-embedded (FFPE) samples of non-diseased human kidney (Figure 1A-D). The 8D6 clone stained interstitial cells (Figure 1A and Supplemental Figure 2), whereas in a serial section S5/1 primarily stained a subset of tubules (Figure 1B). The P12/1 clone (Figure 1C) recognized the same tubules as S5/1 (Figure 1D) in serial section, although the P12/1 staining was weaker. All three clones stained occasional glomerular cells (arrows, Figure 1A-D) and S5/1 occasional interstitial cells, whereas tubular staining was never observed for 8D6. The staining pattern for each antibody was consistent across 4 kidney donors in total, and not observed in the isotype controls (Figure 1E and 1F).

**Figure 1.**
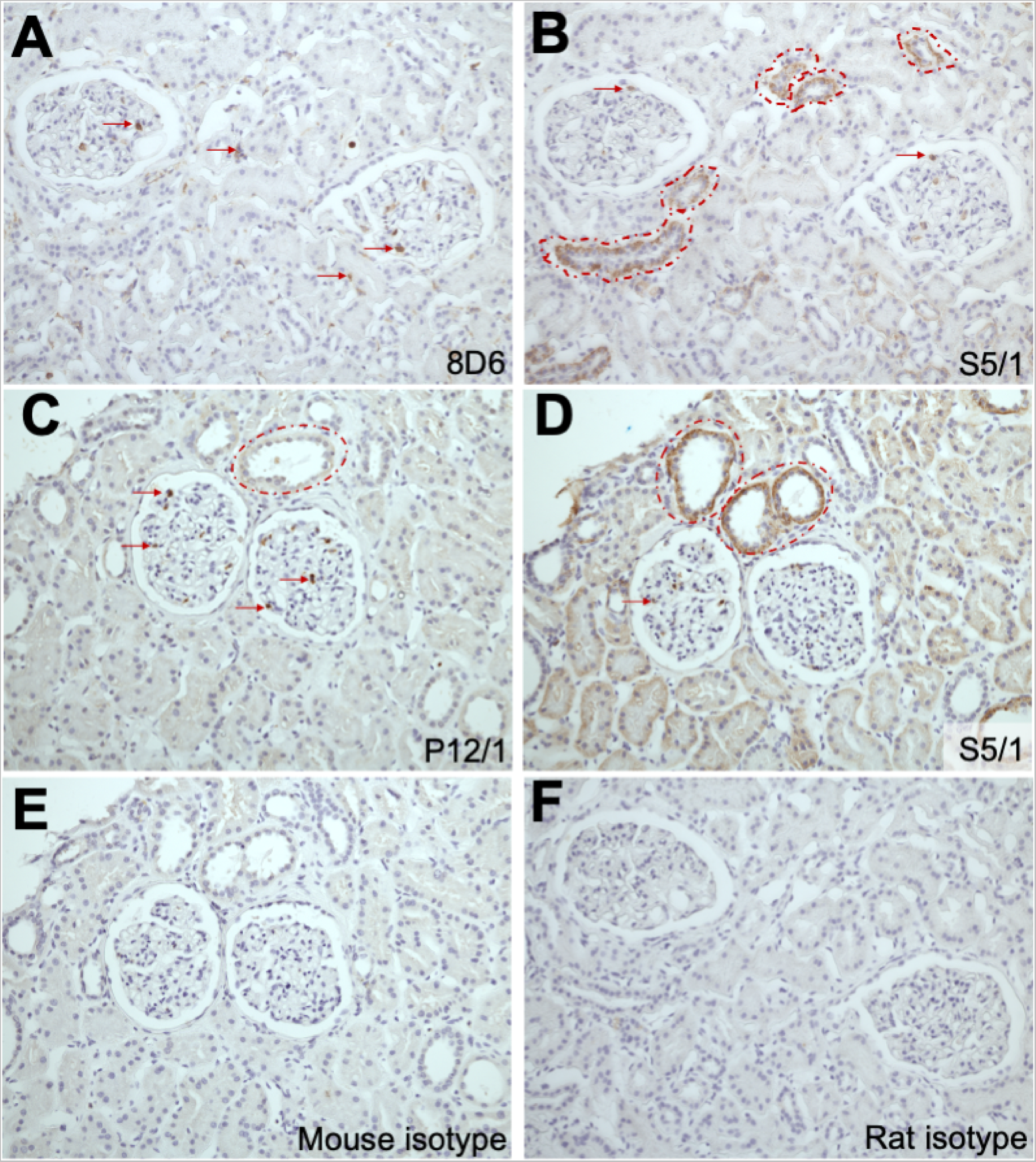
C5aR was expressed in non-diseased human kidney by immunohistochemistry. Representative sections from FFPE human kidney tissue were stained with 3 different C5aR monoclonal antibodies: 8D6 (A), S5/1 (B and D), and P12/1 (C). Red dotted circles and arrows highlight examples of tubules and interstitial (or glomerular) cells, respectively, that are stained brown by the indicated antibody. (E) Section stained with mouse IgG2a antibody as an isotype control for nonspecific binding for S5/1 and P12/1. (F) Section stained with rat IgG2b antibody as an isotype control for 8D6. (A), (B), and (F) are serial sections, as are (C), (D), and (E). Images were counterstained for cell nuclei and are shown at 20x magnification.

We performed in situ hybridization (ISH) with anti-sense probes against C5aR to determine which antibody expression pattern(s) was consistent with mRNA. C5aR transcript was detected in interstitial cells, tubular cells, and occasional glomerular cells in sections of non-diseased human kidney (Figure 2A). Transcript was visualized as purple dots, with expression level proportional to the number of dots per cell, not the intensity of staining. ISH was also performed with the sense probe for C5aR as a negative control, which detected no transcript as expected (Figure 2B), and PPIB, a widely expressed gene, to confirm RNA quality within the section (Figure 2C). A total of 5 human donor kidney samples were examined for C5aR transcript by ISH, and all showed the same pattern of interstitial and tubular staining (for images from an additional 2 donors, see Supplemental Figure 3A-B and 3G-H, with corresponding positive and negative controls shown in Supplemental Figure 3C-F and 3I-L). C5aR transcript levels were higher in interstitial cells, which had tightly packed clusters of dots, than tubule cells, in which individual dots were visible.

**Figure 2.**
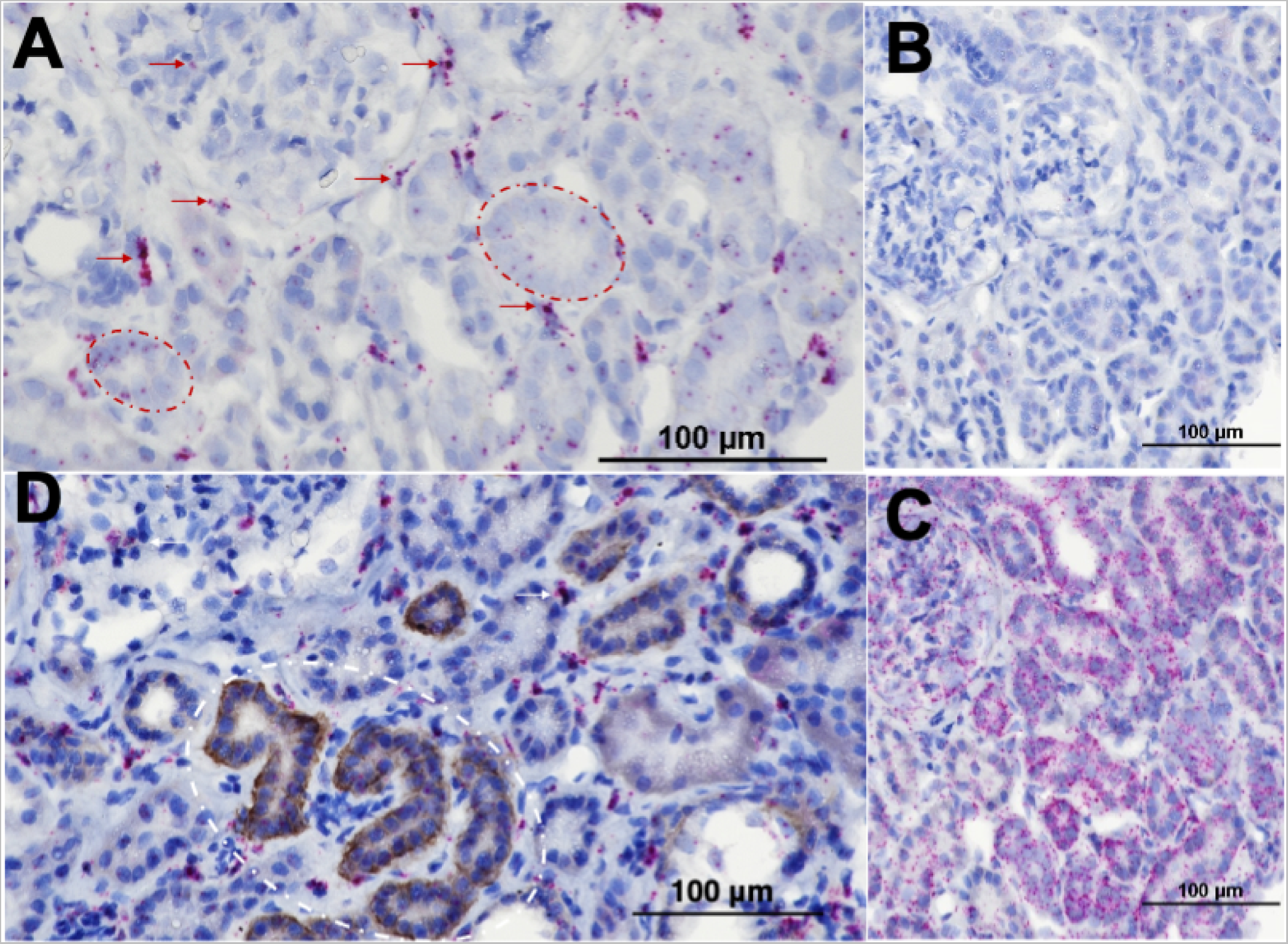
C5aR expression was detected in human kidney tubules and interstitial cells by in situ hybridization (ISH). (A) C5aR transcript was detected in representative human kidney sections with anti-sense oligo probes and visualized as dark purple dots or clusters of dots. Red dotted circles highlight examples of tubule expression; red arrows highlight examples of interstitial and glomerular cell expression. (B) No transcript was detected by the negative control, C5aR sense oligo probes. (C) Expression of the broadly expressed gene PPIB throughout the section confirmed RNA quality in the sample. (D) A dual ISH/IHC method using the S5/1 antibody allowed simultaneous detection of transcript (purple dots) and protein (brown stain) in the same section. White dotted circles highlight examples of tubules and arrows highlight examples of interstitial or glomerular cells positive for both antibody and mRNA expression. All images were counterstained for cell nuclei.

To determine if the ISH and IHC methods detected the same tubules, a dual method was developed with the S5/1 monoclonal antibody. Both C5aR transcript and protein were observed in the same tubules, as highlighted by the dotted circle (Figure 2D). The interstitial and glomerular cell expression observed in ISH was still present in the dual method, and while some of these cells are recognized by both ISH and S5/1 (arrows, Figure 2D), the majority of ISH positive interstitial cells are not stained by S5/1.

Tubule cell subsets express unique proteins that can be used as markers, and dual ISH/IHC methods were developed with antibodies recognizing two of these markers, CALB1 for distal tubules and SLC13A3 for proximal. C5aR transcript was detected in distal tubules (Figure 3A), proximal tubules (Figure 3B), as well as in tubular regions that did not express either marker (Figure 3C and 3D).

**Figure 3.**
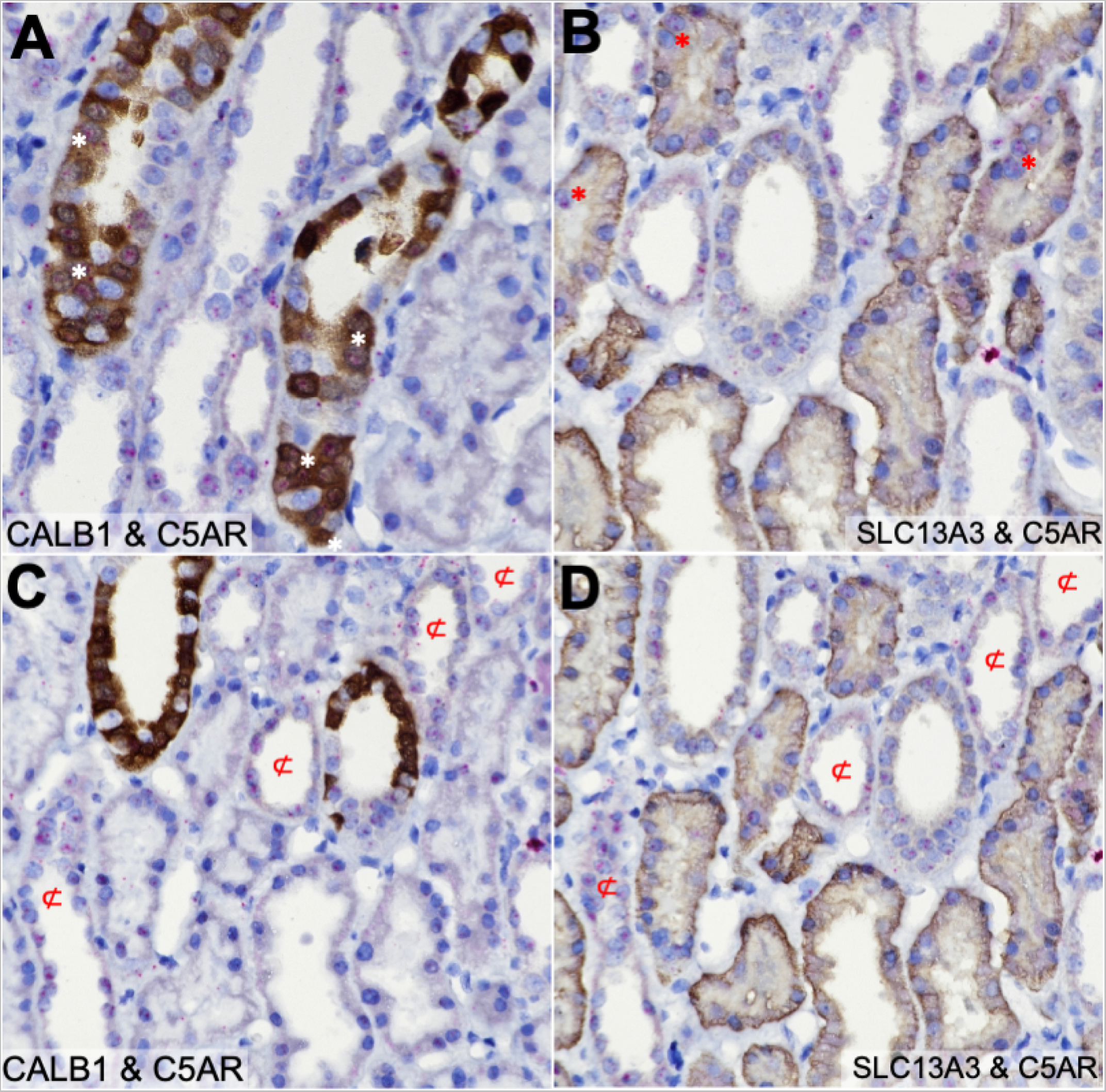
C5aR was expressed in proximal, distal and other human kidney tubule cell subsets. C5aR transcript (purple dots) was detected by ISH (A-D) in non-diseased human kidney FFPE sections that were dual stained by IHC with antibodies for the distal tubule-specific marker CALB1 (brown, A and C) or proximal tubule-specific marker SLC13A3 (brown, B and D). (A, B) Examples of tubule cells expressing both the specified tubule marker and C5aR are highlighted by “*”. (C, D) Tubule regions positive for C5aR transcript but neither tubule marker by serial section overlay are indicated by “⊄”. All images were counterstained for cell nuclei and are shown at 40x magnification.

Dual C5aR ISH/IHC with CD68 demonstrated that the interstitial and glomerular cells expressing C5aR were macrophages (Figure 4A and B). Resident kidney macrophages are dependent on the CSF-1/CSF-1R axis for differentiation, survival, and maintenance (27, 28), and in adults renal macrophages may derive from circulating monocytes as well as precursor cells of embryonic origin (28). Monocytes robustly expressed C5aR across multiple human donors, typically at a higher level than eosinophils and basophils but not neutrophils (Figure 5A). Upon differentiation of monocytes with CSF-1, the expression of C5aR increased further (Figure 5B and C). Monocytes may also be differentiated into macrophages with GM-CSF; C5aR expression is still observed, although at a lower level than CSF-1 macrophages (Figure 5C). C5L2 was expressed by CSF-1-differentiated macrophages, but at best minimally by GM-CSF-differentiated macrophages (Figure 5D).

**Figure 4.**
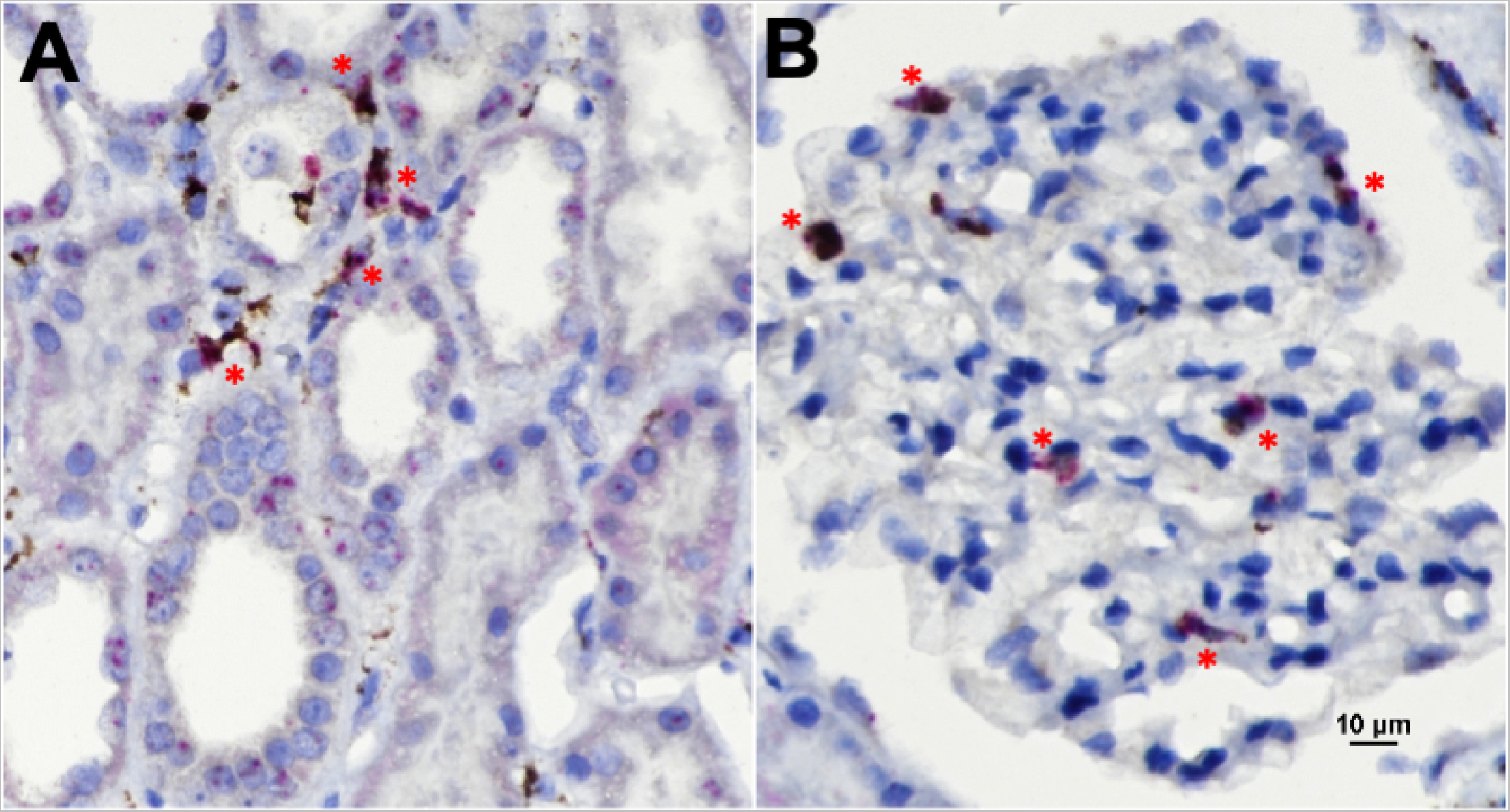
C5aR expression was detected in human kidney resident macrophages and monocyte-derived macrophages. Representative FFPE sections from a non-diseased human kidney donor were dual stained for C5aR by ISH (purple dots) and by IHC with antibodies against the macrophage-specific marker CD68 (brown). (A) C5aR-expressing interstitial cells were CD68 positive macrophages (asterisks). (B) The glomerular cells expressing C5aR were also CD68-positive macrophages (asterisks). All images were counterstained for cell nuclei and the image in A is at 40x magnification.

**Figure 5.**
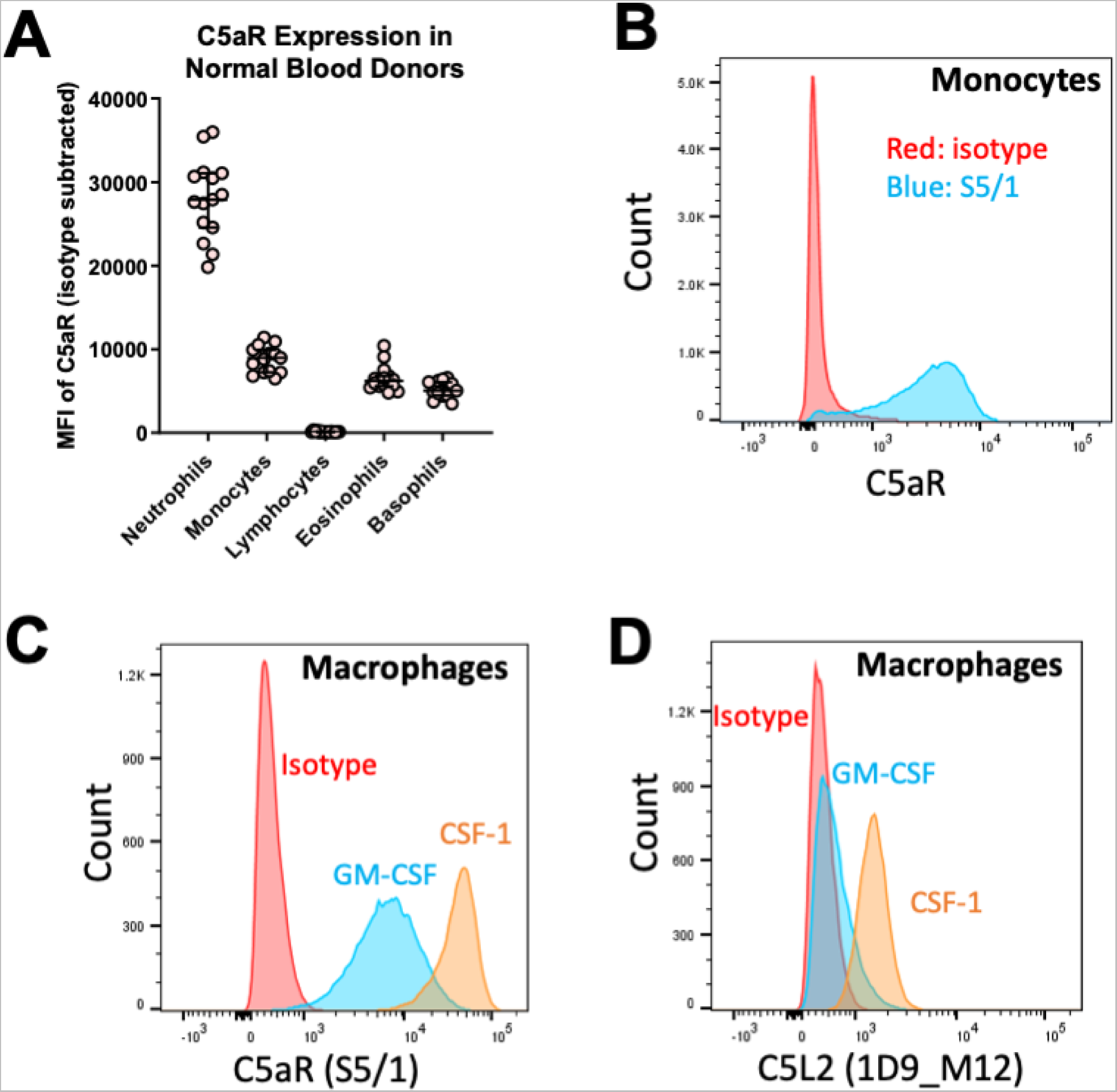
C5aR expression was detected in human leukocytes and monocyte-derived macrophages by flow cytometry. (A) Using PE-labelled C5aR antibody S5/1, the mean fluorescence intensity (MFI) of C5aR in human leukocyte populations is shown for 15 normal human blood donors. Each point on the graph represents the MFI for a single donor, and the median and interquartile range are indicated for each cell type. The cell types were gated as follows: neutrophils, SSC-A^HI^CD16+; monocytes, CD45+CD14+; eosinophils, CD16-CCR3+Siglec8+; basophils, SSC^LO^CD45^DIM^HLADR-CD123+. (B) The histogram shows the distribution of PE-labelled C5aR antibody S5/1 (blue) or isotype (red) stained monocytes isolated from a single representative blood donor. (C) Monocytes from the donor in B were differentiated into macrophages with CSF-1 (orange) or GM-CSF cytokines (blue), then stained with PE-labelled C5aR antibody S5/1 or isotype (red) and analyzed by flow cytometry. (D) An aliquot of the same populations as in C were stained with C5L2 antibody 1D9_M12 (CSF-1 orange and GM-CSF blue) or isotype (red). The histograms in C and D are representative of 3 donors.

### C5L2 kidney expression was detected in a subset of C5aR-positive cells

We investigated the expression of the other C5a receptor, C5L2 (GPR77 or C5aR2), in normal human kidneys using the same ISH and IHC techniques as for C5aR. None of the commercially available C5L2 antibodies demonstrated sufficient specificity and sensitivity for IHC under standard epitope retrieval conditions, thus expression was determined solely by ISH. Kidney C5L2 transcript was detected in interstitial cells and tubule cells at similar levels (Figure 6A), with no more than 1-3 dots per cell. The pattern was compared with C5aR in serial section (Figure 6B), and cells present in both sections that were positive for C5L2 were also positive for C5aR (indicated by asterisks in Figure 6A and B). C5aR transcript was detected in more cells than C5L2, and unlike C5L2 showed higher expression in interstitial cells than in tubule cells (compare Figure 6A to 6B).

**Figure 6.**
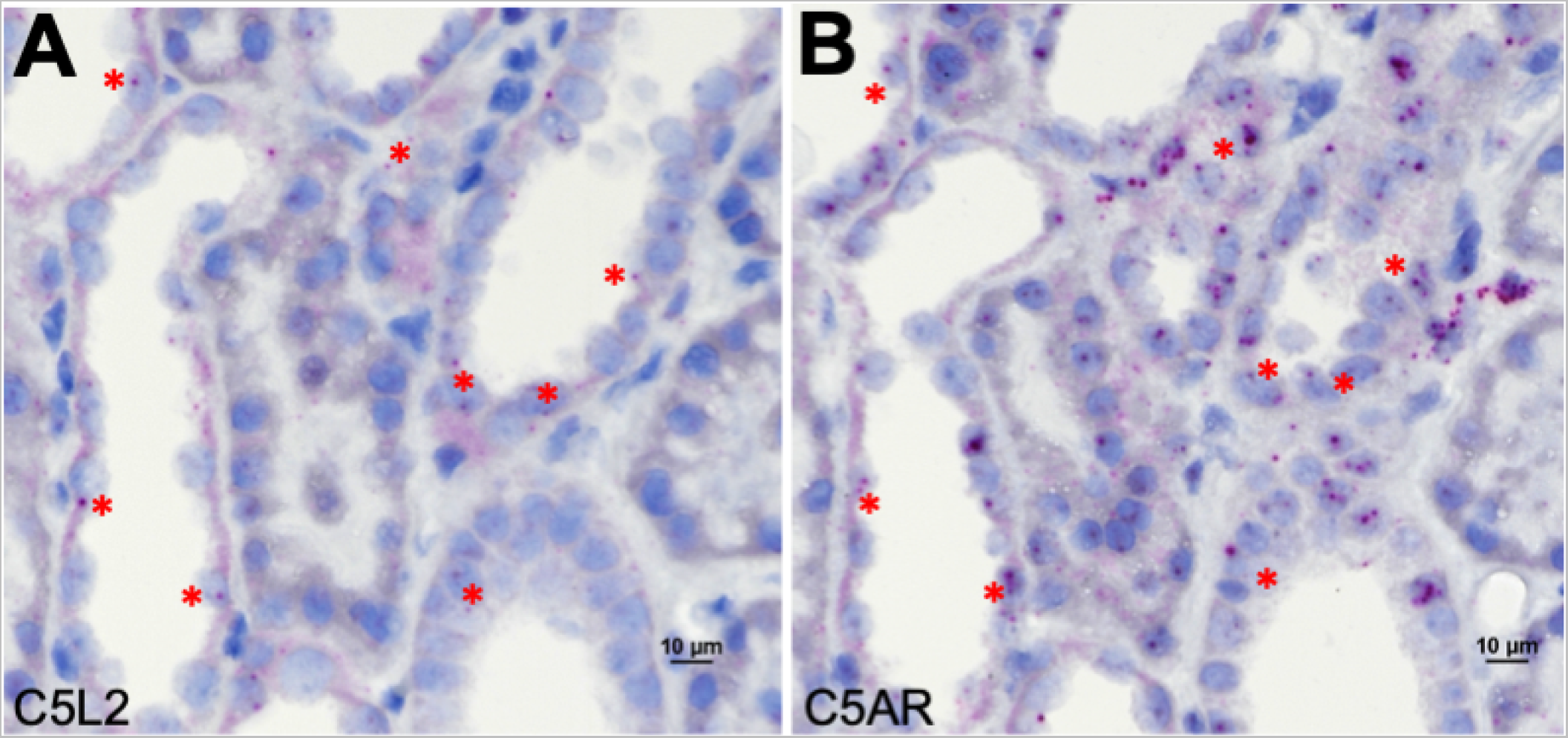
C5L2 transcript was expressed in tubular and interstitial cells positive for C5aR in non-diseased human kidney. Expression of C5L2 (A) and C5aR (B) was detected by ISH in serial kidney sections. Cells present in both sections that express C5aR and C5L2 are indicated by *. Images were counterstained for cell nuclei.

### C5aR was expressed in human C5aR knock-in mouse kidney tubules, interstitial cells, and fibrotic regions

C5aR expression was examined in a C57BL/6 strain in which the mouse C5aR coding sequence was replaced by human C5aR (7). Human C5aR inhibitors such as avacopan do not cross react with mouse C5aR, thus to study disease-modulating effects of avacopan the knock-in strain has been used for translational studies (7), but the expression of human C5aR has not been established in this model system. Human C5aR transcript was detected in both tubules and interstitial cells by ISH (Figure 7A and B), although the transcript level (dots) per cell was lower than in human kidney. Positive control PPIB probes detected uniform transcript levels within sections (Supplemental Figure 4A), indicating RNA was well preserved in the FFPE samples, and the negative control probe showed a complete absence of background transcript detection (Supplemental Figure 4B). The rat monoclonal antibody 8D6 was the only IHC-validated C5aR antibody suitable for mouse, because the antibody backbone was derived from rat and not mouse sequences like S5/1 and P12/1. 8D6 detected human C5aR in the knock-in mouse bone marrow granulocytes by IHC (Supplemental Figure 4C and D) and showed no reactivity against wild type C57BL/6 bone marrow cells (Supplemental Figure 4E and F). No C5aR protein was detected in the knock-in kidney sections stained with 8D6 (Figure 7C), suggesting either C5aR protein level was below the detection limit for 8D6 or that C5aR was expressed as transcript only.

**Figure 7.**
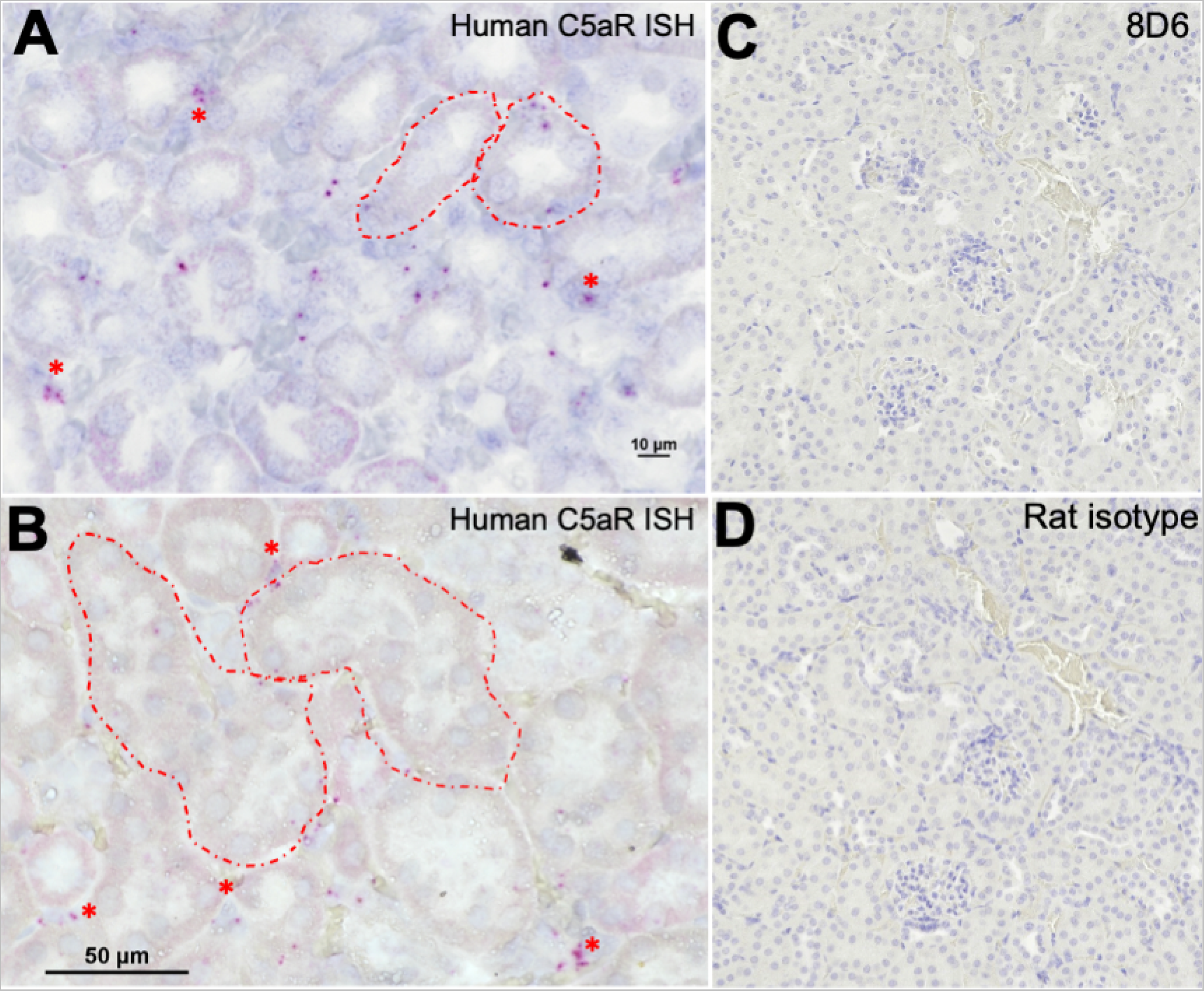
C5aR was expressed in human C5aR knock-in mouse kidney. C5aR expression was examined in kidneys from a human C5aR knock-in mouse line by ISH (A, B) or by IHC with the rat anti-human C5aR monoclonal antibody 8D6 (C). (A, B) Images of two representative fields show that C5aR transcript (purple dots) was detected in tubules and interstitial cells, with examples of tubules highlighted by dotted circles and interstitial cells by asterisks. (C, D) C5aR protein expression was not detected by IHC with the 8D6 antibody (C), which had no staining (brown) in any field, indistinguishable from the isotype control (D). All sections were counterstained for nuclei; C and D are shown at 20x magnification.

Human C5aR expression was then examined in the 5/6 nephrectomy model of chronic kidney disease (29). After surgical removal of kidney tissues, the animals recovered for 6 weeks, at which time the kidney remnants were removed, processed into FFPE blocks, sectioned, and stained with trichrome to identify fibrotic regions of injury (Figure 8A). ISH revealed high levels of C5aR transcript in infiltrating cells within and adjacent to fibrotic regions of nephrectomized kidney (Figure 8B). Consistent with the ISH expression pattern, 8D6 also detected C5aR on numerous interstitial cells in the fibrotic regions of nephrectomized kidney (Figure 8C). In the uninjured areas of the kidneys, 8D6 detection of C5aR decreased, similar to normal mouse kidney (Supplemental Figure 5).

**Figure 8.**
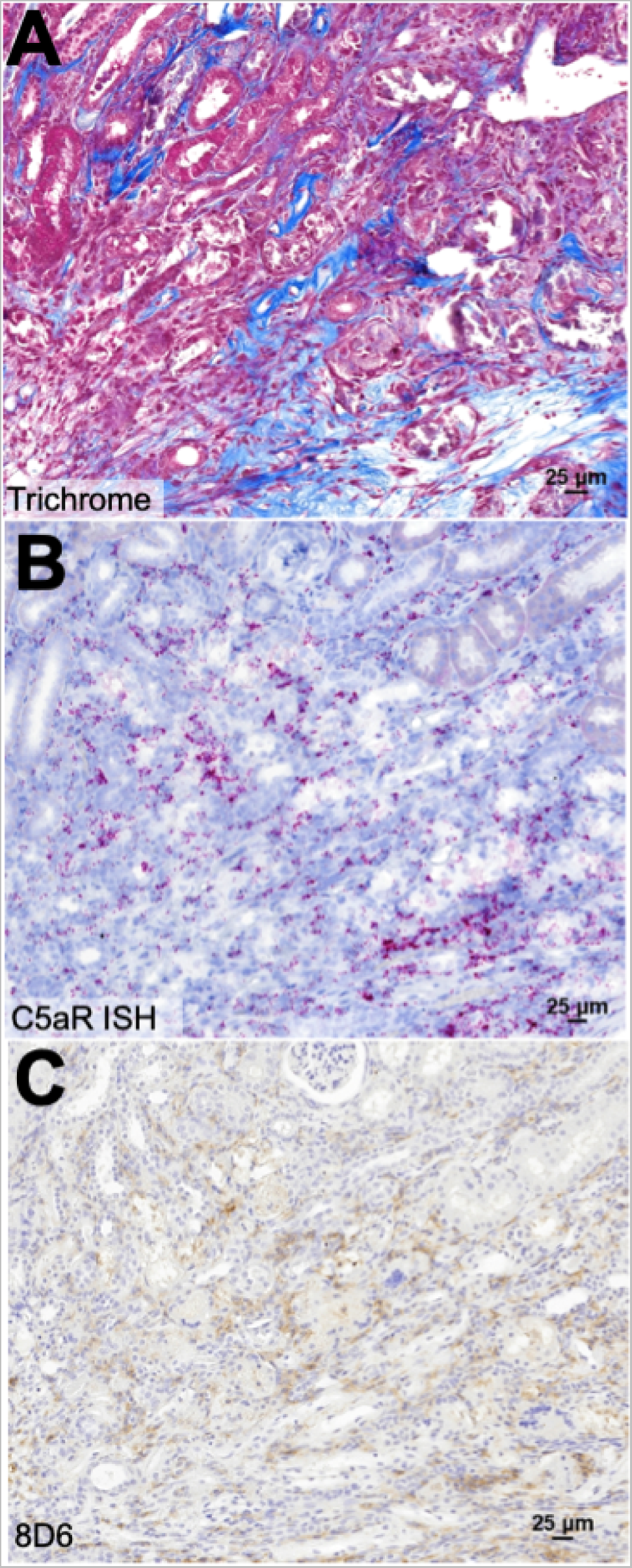
C5aR was expressed in the fibrotic areas of a chronic kidney disease model in human C5aR knock-in mice. (A) An FFPE section of a representative kidney isolated from the human C5aR knock-in mice that underwent 5/6 nephrectomy and were stained with trichrome to reveal regions of fibrosis (blue color). (B) C5aR transcript was detected by ISH (purple dots) in the fibrotic area of the kidney. (C) C5aR protein was also detected by IHC with the antibody 8D6 in the fibrotic area. The images in A, B, and C are all from the same kidney region; B and C are counterstained for cell nuclei.

### Myeloid cell C5aR induced the release of tissue remodeling factors, recruitment of immune cells, and promoted cell survival

The expression of C5aR in kidney resident macrophages and regions of fibrotic kidney injury in a mouse model suggested that C5aR may contribute to inflammation-mediated tissue remodeling. Myeloid cells, including both macrophages and neutrophils, have been implicated in innate immune responses to kidney injury, repair, and fibrosis, and this is an area of active investigation in the field (6, 30, 31).

To model resident and infiltrating kidney macrophages in vitro, primary monocytes from 2 normal donors were differentiated using CSF-1 (28) and treated with C5a, in the presence or absence of the C5aR antagonist avacopan. For an initial survey of the C5aR-dependent macrophage responses, the levels of 21 inflammatory, chemokine, and remodeling factors were assessed (complete list in Methods) using a single high dose of C5a (100 nM). A subset of the secreted protein panel was induced: MMP-3, MMP-8, FAP-α, PDGF-AA, CCL3, CCL7, CCL4, and CCL20 (Figure 9A). F3 (Tissue Factor/CD142), which is a transmembrane protein that initiates the extrinsic coagulation cascade (32), was also upregulated by C5a on the macrophage cell surface (Figure 9B). The addition of avacopan attenuated the responses to C5a, indicating the upregulation of these secreted factors was dependent upon C5aR and not C5L2 activation (Figure 9A and B).

**Figure 9.**
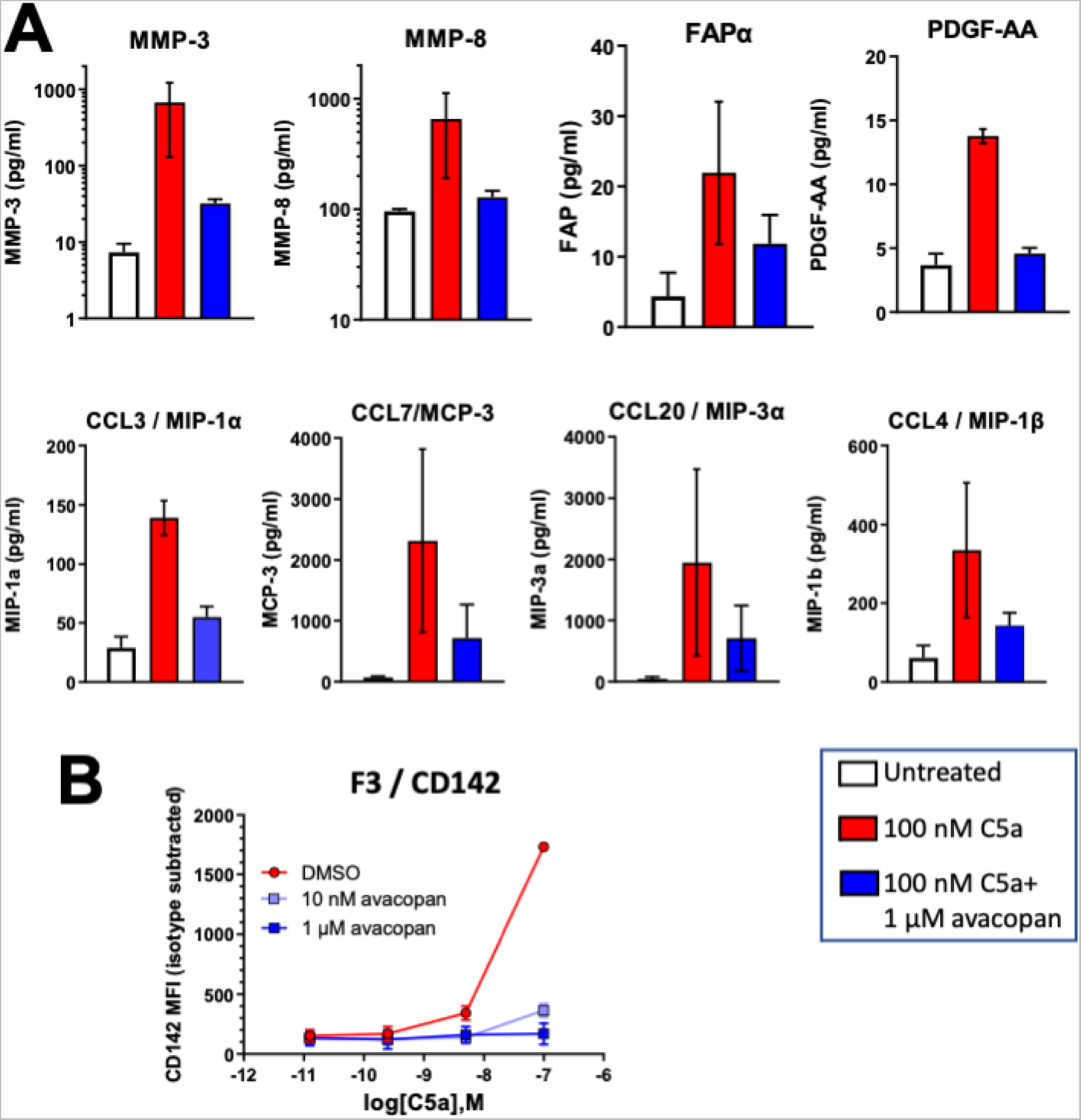
Activation of C5aR on macrophages induced the release of tissue remodeling and immune cell recruitment factors. (A) CSF-1 differentiated macrophages secreted chemokines and tissue remodeling factors in response to 72 hr treatment with 100 nM C5a (red bars). Co-treatment with 1 µM avacopan (blue bars) reduced the response to C5a. (B) C5a dose-dependently induced tissue factor on CSF-1 differentiated macrophages (red line), which was blocked by co-treatment with avacopan (blue lines). The error bars represent range in A and standard deviation in B.

CSF-1 differentiated macrophages from 3 additional normal donors were treated with 100 nM C5a with or without 1 µM avacopan for 24 hr, and the cells were harvested for bulk RNA sequencing. The avacopan-sensitive effects of C5a on macrophage mRNA dynamics were consistent with the secreted protein responses. In addition to MMP-3 and −8, the levels of MMP-10, MMP-1, MMP-19, and the elastase MMP-12, were induced, along with several other factors associated with kidney fibrosis (Supplemental Figure 6A), including but not limited to fibronectin, CD73 (33), Osteopontin (34–36), coagulation factor F5, and the hyaluronidase CEMIP (37, 38). Ligands for the neutrophil chemokine receptors CXCR1 and CXCR2 were also induced by C5aR (Supplemental Figure 6B).

Upon recruitment to injured or diseased kidneys, immune cell activity may be restricted by their short life spans. For myeloid cells, complement may contribute to survival as well as recruitment (39), thus the effect of C5a on apoptosis was examined. Neutrophils were assessed rather than macrophages, since for the latter CSF-1 promotes survival and may complicate the interpretation of in vitro apoptosis. C5a reduced neutrophil apoptosis in response to Fas pathway activation (Figure 10A) and to hydrogen peroxide treatment (Figure 10B) with an IC_50_ of 1 nM C5a, which is comparable in potency to C5a induction of migration (12, 40). Avacopan treatment dose-dependently eliminated the effect of C5a, restoring the apoptotic induction by both pathways (Figure 10C and D). The survival effect of C5a was also abolished by wortmannin (Supplemental Figure 7), suggesting C5aR acts via the PI3 kinase/Akt pathway to mediate blockade of caspase activation and promote survival.

**Figure 10.**
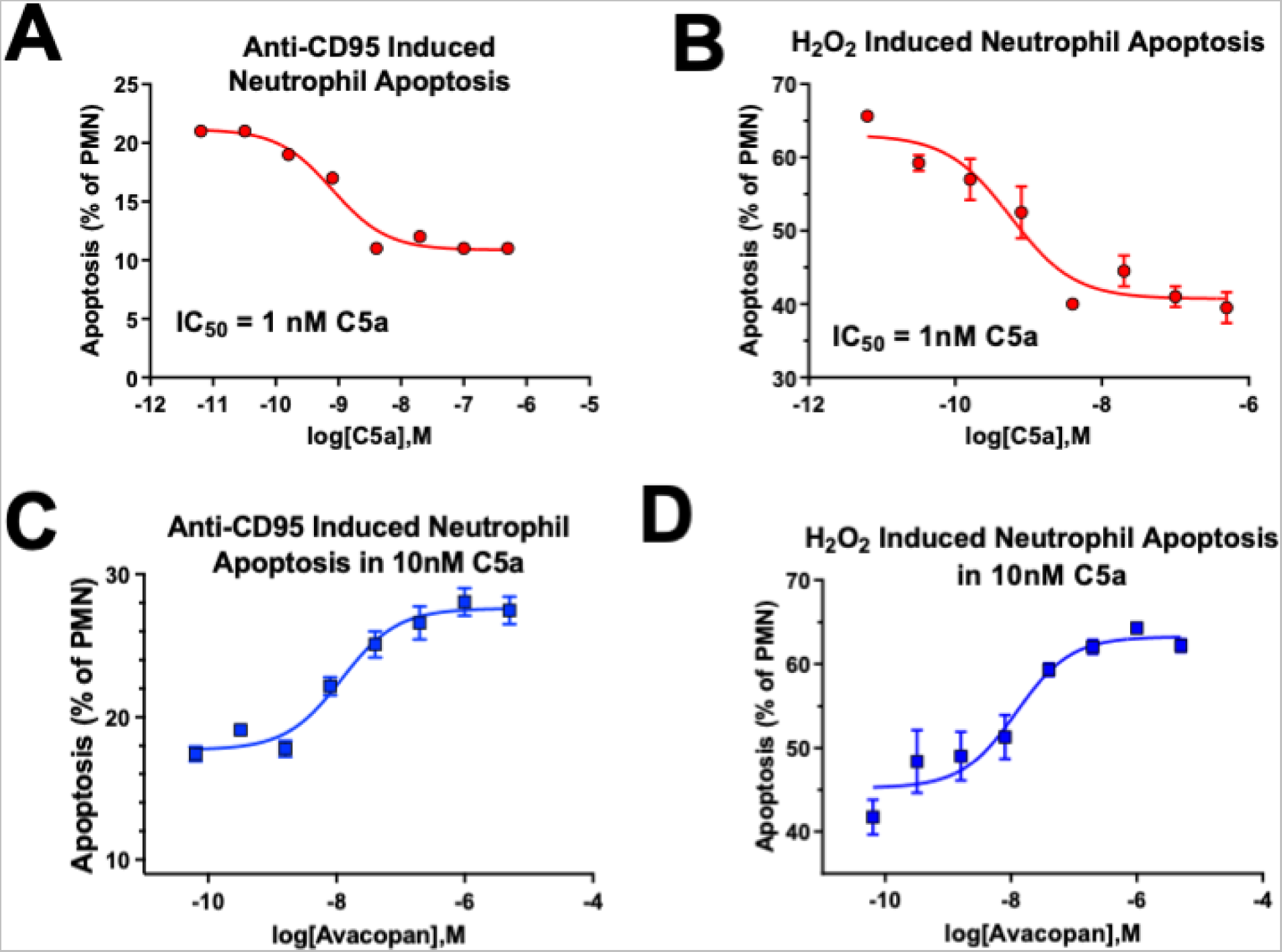
Activation of C5aR on neutrophils inhibited neutrophil apoptosis. C5a dose-dependently increased neutrophil survival in vitro in the presence of apoptosis inducing stimuli anti-CD95 (A) or H_2_O_2_ (B). Based on both dose curves, 10 nM was selected as the C5a dose for assessing avacopan activity. Avacopan treatment restored anti-CD95-induced apoptosis in 10 nM C5a-treated neutrophils induced by anti-CD95 (C) or H_2_O_2_ (D). Error bars represent standard deviation.

## DISCUSSION

### C5a receptor expression in kidney tubule cells and resident macrophages

To ascertain the normal kidney distribution of C5aR it was necessary to reconcile the contrasting results within the literature. As a first step we replicated previous IHC studies in human kidney demonstrating that C5aR monoclonal antibodies S5/1 and P12/1 stained primarily tubules, whereas 8D6 stained only macrophages (14, 15). Protein and mRNA expression levels have been shown to correlate for differentially expressed genes (41), suggesting that the C5aR protein expression should be identical to or a subset of the C5aR mRNA distribution.

By ISH we detected the highest C5aR transcript in macrophages, consistent with 8D6 expression, and lower levels across tubule subsets. The high tubule and minimal macrophage staining intensities of S5/1 and P12/1 were not consistent with ISH, or with the fact that both antibodies robustly recognized myeloid cells by IHC in our hands and macrophages in other organs (20, 24). Explanations for this discrepancy include antibody polyreactivity against a tubule epitope (15) and differential recognition of cell type-specific C5aR post-translational modifications. Consistent with these explanations, the P12/1 and S5/1 antibodies have epitopes distinct from 8D6 within the N-terminus, and both were raised using a peptide immunogen, whereas cell-expressed C5aR was used for 8D6 (15, 42).

Our results strongly support incorporating a robust ISH method in C5aR tissue expression studies. The 8D6 antibody is the most selective reagent for C5aR IHC in our hands, and has the added benefit of suitability for mouse tissue. Investigation of C5aR expression with these methods in patient kidneys and other organs will provide insight into the mechanism of avacopan in ANCA vasculitis (10), as well as in other diseases such as lupus with pathology linked to fibrosis or complement system activation.

In contrast, C5L2 protein expression could not be assessed by IHC, because no commercial antibody demonstrated sufficient specificity under standard epitope retrieval conditions, including previously published polyclonal reagents (20). C5L2 transcript was detected in a subset of the C5aR positive cells by ISH, and was co-expressed in in vitro CSF-1 differentiated macrophages by flow cytometry. C5L2 expression was consistently lower than C5aR in every cell type examined, but without reference standards an absolute comparison of expression cannot be performed. Since the presence or absence of C5L2 may significantly affect C5aR activity (8), its expression pattern provides critical context for C5aR and should always be assessed in parallel.

### C5aR functional relevance to kidney fibrosis

The human C5aR KI mouse strain expressed C5aR by ISH in kidney tubules and interstitial cells, a pattern similar to human. When the mice were subjected to 5/6 nephrectomy, elevated C5aR was detected specifically in regions of kidney fibrosis by ISH and also by IHC with 8D6. A qualitative increase in both C5aR positive infiltrating cells and cellular expression was observed, compared to surrounding tissue. The infiltrating C5aR positive cells appeared to be largely macrophages, although a dual method was not used to confirm this.

These observations are consistent with an elegant series of experiments in the folic acid model of kidney fibrosis with a floxed C5aR mouse line (6), which demonstrated that macrophages were the major C5aR- positive interstitial cells in kidney and that folic acid treatment increased both the number and C5aR expression level of these macrophages. Furthermore, deleting C5aR specifically from macrophages significantly reduced folic acid-induced fibrosis, suggesting C5aR mediates macrophage contributions to the development of fibrosis in this model.

To identify contributions of myeloid C5aR to human kidney fibrosis, we examined C5aR activation by C5a and inhibition by avacopan in CSF-1 differentiated macrophages. In vitro studies with macrophages are limited by the lack of kidney-specific factors that modulate differentiation and function, but both populations share dependence on CSF-1 and derivation from monocytes. These studies revealed that C5aR induced the secretion of chemokine ligands capable of recruiting neutrophils, monocytes, and T cells, and upregulated a number of factors involved in tissue remodeling that may contribute to kidney fibrosis, including F3, multiple MMPs, PDGF-AA, and FAP. Notably, F3 is a master regulator of the extrinsic coagulation system that is thought to exacerbate renal injury by enhancing renal inflammation and fibrosis (32). Some or all of these factors could thus contribute to ANCA-associated vasculitis pathology in the kidney, as well as other organs. For example, the C5aR-mediated upregulation of the elastase MMP12 by in vitro macrophages may contribute to the erosion of elastic cartilage observed in some ANCA patients, resulting in damage to the kidney but also the lung, ear, and nose (43).

We were unable to examine C5aR function in kidney tubule cells in vitro, as no primary cells were available that expressed C5aR. Tubular C5aR may have a role distinct from macrophage/myeloid C5aR in kidney pathology. For example, C5aR deficiency has been reported to protect against pathogenic inflammation, fibrosis and bacterial load in a model of chronic pyelonephritis (44), thus tubular C5aR could play a role in innate responses to ureter or bladder infections.

The functional studies for C5aR in macrophages provide a basis for generating a C5aR activation signal, which awaits further validation in the exogenous C5a (45) and ANCA vasculitis (7) models using multi-dimensional spatial imaging or similar approaches. Tissue- and cell type-specific C5aR function may then be examined in response to avacopan treatment in both models and human disease. This would also provide an opportunity to further map tubular cell types expressing C5aR and, in concert with targeted deletions, to explore tubule-specific functions of C5aR.

## ACKNOWLEDGMENTS

We would like to thank Israel Charo for advice and feedback during the course of these studies.

## AUTHOR CONTRIBUTIONS

All authors conceived and designed research, C.D., N.Z., and L.E. performed experiments, C.D., N.Z., and

K.S. analyzed data, interpreted results of experiments, and prepared figures. K.S. drafted the manuscript and prepared the final revision. All authors edited the manuscript and approved final version of manuscript.

